# Fenretinide inhibits obesity and fatty liver disease but induces Smpd3 to increase serum ceramides and worsen atherosclerosis in LDLR-/- mice

**DOI:** 10.1101/2022.07.22.500933

**Authors:** Dawn Thompson, Shehroz Mahmood, Nicola Morrice, Sarah Kamli-Salino, Ruta Dekeryte, Philip Hoffmann, Mary K. Doherty, Philip D. Whitfield, Mirela Delibegović, Nimesh Mody

## Abstract

Fenretinide is a synthetic retinoid that can prevent obesity and improve insulin sensitivity in mice by directly altering retinol/retinoic acid homeostasis and inhibiting excess ceramide biosynthesis. We determined the effects of Fenretinide on LDLR^-/-^ mice fed high-fat/high-cholesterol diet +/- Fenretinide, a model of atherosclerosis and non-alcoholic fatty liver disease (NAFLD). Fenretinide prevented obesity, improved insulin sensitivity and completely inhibited hepatic triglyceride accumulation, ballooning and steatosis. Moreover, Fenretinide decreased the expression of hepatic genes driving NAFLD, inflammation and fibrosis e.g. *Hsd17b13, Cd68* and *Col1a1*. The mechanisms of Fenretinide’s beneficial effects in association with decreased adiposity were mediated by inhibition of ceramide synthesis, via hepatic DES1 protein, leading to increased dihydroceramide precursors. However, Fenretinide treatment in LDLR^-/-^ mice enhanced circulating triglycerides and worsened aortic plaque formation. Interestingly, Fenretinide led to a 4-fold increase in hepatic sphingomyelinase *Smpd3* expression, via a retinoic acid-mediated mechanism and a further increase in circulating ceramide levels, linking induction of ceramide generation via sphingomyelin hydrolysis to a novel mechanism of increased atherosclerosis. Thus, despite beneficial metabolic effects, Fenretinide treatment may under certain circumstances enhance the development of atherosclerosis. However, targeting both DES1 and Smpd3 may be a novel, more potent therapeutic approach for the treatment of metabolic syndrome.

## INTRODUCTION

Obesity has reached epidemic proportions worldwide and contributes to the pathophysiology of many disease states including type 2 diabetes, cardiovascular disease, and non-alcoholic fatty liver disease (NAFLD) collectively referred to as metabolic syndrome. Indeed, many of these diseases have overlapping secondary pathologies and are risk factors for the development of further complications. For example, defective insulin signalling associated with dyslipidaemia and chronic low-grade inflammation results in increased risk of developing type-2 diabetes, NAFLD and atherosclerosis (1–3). Specifically, NAFLD, is now the most common liver disease in the Western world, characterised by an accumulation of triglycerides that can develop from simple steatosis to non-alcoholic steatohepatitis (NASH) and progress to cirrhosis and hepatocellular carcinoma (4,5). Although there are therapeutics available for type 2 diabetes and CVD, there are no approved treatments currently available for NAFLD other than dietary and lifestyle intervention/changes. Since it is becoming increasingly common for patients to present with multiple overlapping co-morbidities, there is an urgent need for new therapeutics targeting the mechanistic causes of these diseases to halt the predicted rise in cases.

Increased lipotoxicity and accumulation of the bioactive mediator ceramide has been attributed to be a major player in the progression of obesity-associated metabolic diseases (6). Ceramide belongs to the sphingolipid class of lipid mediators, the generation of which is tightly controlled by a series of enzymes through either *de novo* synthesis, sphingomyelin hydrolysis or through the salvage pathway. Indeed, there have been numerous studies reporting high fat diet feeding leading to increased *de novo* sphingolipid synthesis resulting in the accumulation of ceramide in several tissues such as liver, adipose tissue, skeletal muscle and the heart (7). Nonhuman primates fed a western diet to induce obesity and type 2 diabetes exhibited increased circulating ceramides (8,9). Circulating ceramides are also considered important risk factors for cardiovascular disease (10). Mechanistically, excess ceramide has been shown to lead to defective insulin signalling due to an impairment of the downstream effector cascades such as activation of Akt or GSK3B (11,7,12). Therefore, inhibiting ceramide accumulation is an attractive target for manipulation.

Fenretinide (FEN, also known as N-(4-hydroxyphenyl)retinamide or 4-HPR)) is a synthetic derivative of retinoic acid and has been investigated as a potential therapeutic for metabolic syndrome. In previous studies FEN treatment led to decreased weight gain and adiposity and improved glucose homeostasis and insulin sensitivity in association with decreased accumulation of hepatic triglycerides in high-fat diet fed mice (13–18). The mechanism of FEN action has been attributed to alterations in retinol homeostasis and retinoic acid signalling and prevention of lipotoxicity by directly inhibiting the elevation of several ceramide species both *in vitro* (19) and *in vivo*, in adipose (14), liver tissue (17) and skeletal muscle (20). Interestingly, *Smpd3* encodes for a type 2-neutral sphingomyelinase (nSMase2) that has been identified as transcriptionally induced by retinoic acid (21,22). *Smpd3/nSMase2* is key to an alternative ceramide generation pathway (via sphingomyelin hydrolysis) that has recently been linked to atherosclerosis via regulation of serum ceramide levels (23).

There have been several models, both dietary and genetic, used to study the effect of obesity on either type 2 diabetes, atherosclerosis or NAFLD, and some of these also to determine the mechanism of FEN action. Since many of the diseases associated with obesity have overlapping pathologies, we sought to determine if FEN could improve several disorders in LDLR^-/-^ mice, traditionally a model for atherosclerosis with a similar lipoprotein profile to humans (24). On a high-fat/high-cholesterol diet, LDLR^-/-^ mice become obese, develop insulin resistance and accumulate hepatic triglycerides, with inflammation and hepatic fibrosis associated with NAFLD progression to NASH (25,26). Given previous research has shown blocking ceramide biosynthesis is beneficial (6,9), we hypothesized that FEN could be used as a novel intervention for NAFLD/NASH and atherosclerosis, in addition to its beneficial effects of decreased adiposity and improved insulin sensitization, via prevention of excess ceramide accumulation.

## RESEARCH DESIGN AND METHODS

### Animal studies

All animal procedures were performed under a project licence (PPL P94B395E0) approved by the U.K. Home Office under the Animals (Scientific Procedures) Act 1986. Male LDLR^-/-^ mice, aged 4-6 weeks, were purchased from The Jackson Laboratory (supplied by Charles River UK Ltd), male and female ApoE^-/-^ mice were bred in-house (University of Aberdeen). All mice were fed chow diet until 12 weeks of age then placed into three groups and fed control (10% kCal fat D14121001) or high-fat/high-cholesterol diet (HFD, 40% kCal fat plus 1.25% cholesterol, D12108C) for 14 weeks to induce atherogenesis and NAFLD +/− 0.04% Fenretinide (FEN-HFD, D18061502, Research Diets Inc). Mice were maintained at 22–24°C on 12-h light/dark cycle with free access to food/water. At week 14, mice were fasted for 5h and injected intraperitoneally with either saline or insulin (10 mU/g body weight) for 10 mins prior to CO2-induced anaesthesia followed by cervical dislocation. Heart and aortic tissues were collected for histological analysis. Peripheral metabolic tissues (liver, muscle and white adipose tissue (WAT)) were frozen in liquid nitrogen and stored at −80°C until subsequent analysis.

### Glucose and Insulin Tolerance Tests

Mice were fasted for 5h prior to commencement of glucose or insulin tolerance tests (GTT and ITT, respectively). Briefly, baseline glucose levels were sampled from tail blood using glucose meters (AlphaTRAK, Abbott Laboratories, Abbot Park, IL, USA). Subsequently mice were injected intraperitoneally with 20% glucose (w/v) or insulin (0.75mU/g body weight) and blood glucose measured at 15-, 30-, 60- and 90-mins postinjection.

### Body Fat Mass Composition

The body composition of mice was analysed using an Echo MR 3-in-1 scanner (Echo Medical Systems, Houston, TX, USA).

### Immunoblotting

Frozen liver tissues were homogenised in 400μl of ice-cold RIPA buffer (10mM Tris-HCl pH 7.4, 150mM NaCl, 5mM EDTA pH 8.0, 1mM NaF, 0.1% SDS, 1% Triton X-100, 1% Sodium Deoxycholate with freshly added 1mM NaVO_4_ and protease inhibitors) using a PowerGen 125 homogeniser and lysates normalised to 1μg per 1μl. Proteins were separated on a 4-12% Bis-Tris gel by SDS-PAGE and transferred onto nitrocellulose membrane. Membranes were probed for the following; phospho-AKT (Ser 473, cat: 4060), total Akt (cat: 4691), phospho-S6 (Ser 235/236, cat: 4858), total S6 (cat: 2217), phospho-AMPK (Thr 172, cat: 2535)), total AMPK (cat: 5832) and GAPDH (cat: 5174) (all Cell Signaling Technology), or IR β-chain (Santa Cruz Biotechnology). ApoB 48, ApoB 100 (Meridian Life Sciences UK, cat: K23300R) and Vinculin (Cell Signaling Technology, cat: 13901)) were separated on a 6% Tris-Glycine gel. Anti-rabbit and anti-mouse horse radish peroxidase (HRP) conjugated antibodies were from Anaspec. Primary and secondary antibodies were used at 1:1000 and 1:5000 respectively.

### RNA extraction and qPCR

Frozen tissues were lysed in TRIzol reagent (Sigma, U.K.) and RNA isolated using phenol/chloroform extraction according to manufacturer’s instructions. RNA was then synthesized into cDNA (tetrokit, Bioline) and subjected to qPCR analysis using SYBR green and LightCycler 480 (Roche). Gene expression was determined relative to the reference gene□*Nono* or *Ywaz*. Details of primer sequences can be found in Supplemental Table 1.

### Histology

Liver tissues were sectioned and stained to assess steatosis (Haemotoxylin and Eosin (H&E) or fibrosis (picrosirius red). Immediately following cervical dislocation, hearts with attached aortic root were immersed in formalin and stored at 4°C for 24hrs, before being transferred to PBS until further analysis. Hearts were bisected to remove the lower ventricles, frozen in OCT and subsequently sectioned at 5μm intervals until the aortic sinus was reached. Sections were mounted and stained with oil red O to assess plaque formation. The descending aorta was prepared for *en face* staining. Briefly aortas were trimmed of perivascular adipose tissue, cut longitudinally, and stained with Sudan IV to assess plaque formation. Images were captured using a light microscope and plaque formation quantified using Image J software. Plaque formation in aortic root sections was represented as total area whereas for *en face* staining percentage plaque area was used.

### Liver Triglyceride Assay

50-100mg of frozen liver tissue was homogenised in 1ml of PBS and frozen in liquid nitrogen to enable further cell lysis. Samples were thawed, centrifuged briefly (15s at 7500rpm) and the supernatant (including the lipid layer on top) transferred to a fresh tube. Total triglycerides were measured in homogenates according to manufacturer’s instructions (Sigma, cat: MAK266).

### Serum Analysis

Blood was collected during terminal procedures after fasting (5hrs) and spun to isolate serum, then stored at −80°C. Serum samples were subsequently analysed for total cholesterol and triglycerides (Sigma, cat: MAK043 and MAK266 respectively) or Insulin and Leptin (Crystal Chem, cat 90080 and 90030 respectively) according to manufacturer’s instructions.

### Quantification of liver dihydroceramides and ceramides

Extraction of liver lipids was performed according to the method described by Folch *et al.* (27). Dihydroceramides and ceramides and were isolated by solid phase extraction chromatography using C12:0 dihydroceramide and C17:0 ceramide (Avanti Polar Lipids, Alabaster, Al, USA) as internal standards. Samples were analysed by liquid chromatography-mass spectrometry (LC-MS) using a Thermo Exactive Orbitrap mass spectrometer (Thermo Scientific, Hemel Hempsted, UK) equipped with a heated electrospray ionization (HESI) probe and coupled to a Thermo Accela 1250 ultra-high-pressure liquid chromatography (UHPLC) system. Samples were injected onto a Thermo Hypersil Gold C18 column (2.1 □mm by 100□mm; 1.9□μm) maintained at 50°C. Mobile phase A consisted of water containing 10□mM ammonium formate and 0.1% (vol/vol) formic acid. Mobile phase B consisted of a 90:10 mixture of isopropanol-acetonitrile containing 10 rmM ammonium formate and 0.1% (vol/vol) formic acid. The initial conditions for analysis were 65% mobile phase A, 35% mobile phase B and the percentage of mobile phase B was increased from 35% to 65% over 4□min, followed by 65% to 100% over 15 min, with a hold for 2 min before reequilibration to the starting conditions over 6 min. The flow rate was 400□ulmin and samples were analyzed in positive ion mode. The LC-MS data were processed with Thermo Xcalibur v2.1 (Thermo Scientific) with signals corresponding to the accurate mass-to-charge ratio (m/z) values for dihydroceramide and ceramide molecular species extracted from raw data sets with the mass error set to 5□ppm. Quantification was achieved by relating the peak area of the dihydroceramide and ceramide lipid species to the peak area of their respective internal standard. All values were normalised to the wet weight of liver.

### Statistical Analysis

Data are presented as mean +/± S.E.M. Group sizes were determined by performing a power calculation to lead to an 80% chance of detecting a significant difference *(P* ≤ 0.05). For both *in vivo* and *ex vivo* data, each *n* value corresponds to a single mouse. Statistical analyses were performed by using one-way or two-way ANOVA followed by Bonferroni multiple-comparison tests to compare the means of three or more groups. Variances were similar between groups. In all figures, */^#^p<0.05, **/##p≤0.01, ***/^###^p<0.001, ****/^####^p≤0.0001. All analyses were performed using GraphPad Prism (GraphPad Software).

### Data and Resource Availability

The data sets generated and analyzed during the current study are available from the corresponding author upon reasonable request. RNA-Seq data referred to here and (17) will be deposited at the Gene Expression Omnibus (GEO) website following publication of manuscript in preparation. Fenretinide used in the current study is available from the corresponding author upon reasonable request and for non-commercial purposes (17).

## RESULTS

### Fenretinide prevents diet-induced obesity and improves insulin sensitivity in LDLR^-/-^ mice fed an atherogenic diet

Male LDLR^-/-^ mice were fed an obesogenic plus atherogenic, high-fat/high-cholesterol diet (HFD) +/− 0.04% FEN (FEN-HFD) or control diet for 14 weeks. All mice gained body weight until about week 8 when HFD mice continued to gain body weight but FEN-HFD mice and control mice body weights reached a similar plateau for the remainder of the study (**Fig. 1A**). This inhibition of body weight gain was due specifically to an inhibition of adiposity in FEN-HFD mice and not due to alterations in lean mass **(Fig. 1B, 1C**). Serum leptin levels were markedly elevated in HFD mice whereas in FEN-HFD mice levels were similar to control mice (**Table 1**).

**Figure 1:**
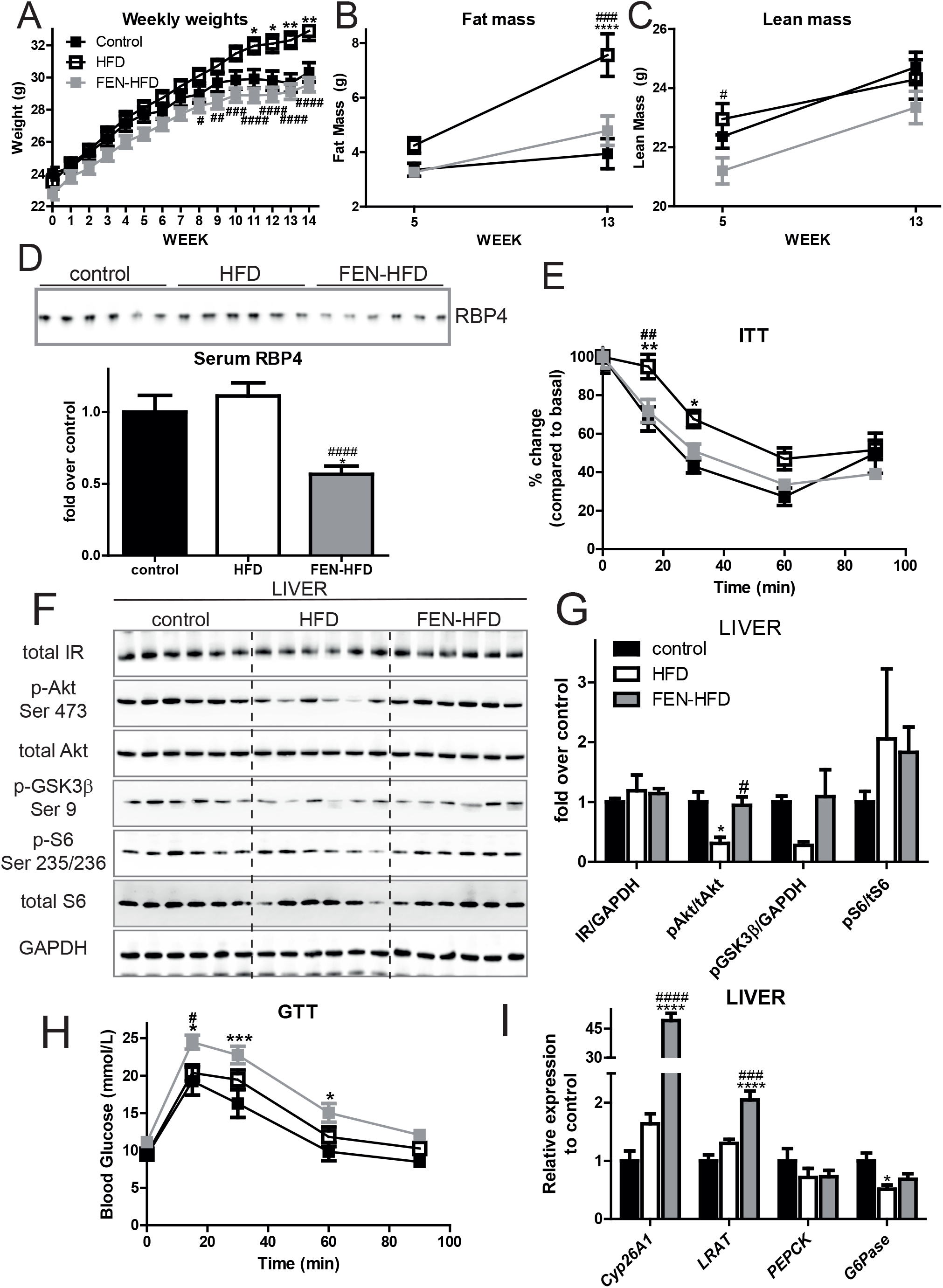
Fenretinide prevents diet-induced obesity and improves insulin sensitivity in the LDLR^-/-^ mice. (**A**) Weekly body weights of LDLR^-/-^ mice fed either control (n=14, 10% kCal fat) or high-fat/high-cholesterol diet +/− 0.04% Fenretinide (HFD, n=18, 40% kCal fat plus 1.25% cholesterol or FEN-HFD, n=18 respectively). Body composition total body fat mass (**B**) and lean mass (**C**) at week 5 and week 13 diet. (**D**) Serum RBP4 levels (n=6 per group). (**E**) Insulin tolerance test (ITT) at week 13 (n=10 per group). At 14 weeks diet, liver tissue western blot analysis (F, *representative image*) and quantification of bands (G, n=10 per group) from LDLR-/- mice injected with insulin (10 mU/g body weight, 10 mins). (**H**) Glucose tolerance tests (GTT) at week 11. (**I**) Hepatic expression of RA/RAR target genes (n=8 per group). Data are represented as mean ± S.E.M. and analysed by either Two-way (**A**-**C, E, H**) or one-way ANOVA (**D, G, I**) followed by Bonferroni multiple comparison t-tests where *p≤0.05, **p≤0.01, ***p≤0.001 and ****p≤0.0001 (control compared to HFD) or #p≤0.05 and ## p≤0.01 and ### p≤0.001 (HFD compared to FEN-HFD).

**Table 1:** Serum and Tissue measurements. Data are represented as mean ± S.E.M. (n=8 per group) and analysed by One-way ANOVA followed by Bonferroni multiple comparison t-tests where **p≤0.01, ***p≤0.001 and ****p≤0.0001 (control compared to HFD) or ## p<0.01 and #### p≤0.0001 (HFD compared to FEN-HFD).

|  | control | HFD | FEN-HFD |
| --- | --- | --- | --- |
| **Serum levels** |  |  |  |
| Basal Glucose Week 11 | 9.32 ± 0.57 | 9.51 ± 0.76 | 11.12 ± 0.51 |
| Basal Glucose Week 12<br>(mmol/L) | 7.65 ± 0.65 | 8.67 ± 0.57 | 9.75 ± 0.56 |
| Insulin (ng/ml) | 1.23 ± 0.37 | 0.82 ± 0.12 | 0.69 ± 0.07 |
| Leptin (ng/ml) | 6.94 ± 3.03 | 15.79 ± 3.75 | 7.56 ± 1.43 |
| Cholesterol (µg) | 5.96 ± 1.00 | 17.21 ± 1.60 **** | 17.19 ± 0.57 **** |
| Triglyceride (mg/ml) | 1.76 ± 0.195 | 4.41 ± 0.43 *** | 7.95 ± 0.60 ****/#### |
| **Tissue levels** |  |  |  |
| Liver triglyceride (µg/mg) | 14.24 ± 1.47 | 35.28 ± 6.34 ** | 13.82 ± 0.97 ## |

As expected, FEN treatment decreased serum RBP4 levels compared to levels in control and HFD LDLR -/- mice (**Fig. 1D**). However, several classic molecular markers of functional white adipose tissue (e.g. *PPARy, GLUT4)* were largely unaltered in LDLR^-/-^ mice fed HFD +/− FEN, although there was a near 50% decrease in white adipose *PEPCK, adiponectin, resistin* and *RBP4* (**Supplemental Fig. 1**). HFD induced physiological insulin resistance (**Fig. 1E**) and decreased acute hepatic insulin signalling to Akt (**Fig. 1F, 1G**). Whereas FEN treatment, resulted in improved insulin sensitivity and rescued hepatic Akt phosphorylation in response to insulin (**Fig. 1F, 1G**).

Despite these markedly beneficial physiological effects (decreased adiposity and improved insulin sensitivity) other parameters of glucose homeostasis were not similarly improved with FEN treatment. Basal serum glucose and serum insulin levels (in the 5-h fasted state) were similar in all three LDLR^-/-^ groups and FEN treatment increased glucose intolerance compared to both HFD and control LDLR^-/-^ mice (**Table 1 and Fig. 1H**). This effect of FEN was not attributable to changes in hepatic *PEPCK*, a known RA/RAR target gene (**Fig. 1I**) despite induction of hepatic *Cyp26A1* and *LRAT*, classic RA/RAR target genes (**Fig. 1I**). In skeletal muscle, acute hepatic insulin signalling to Akt was not altered by HFD +/− FEN. FEN-HFD fed mice had significantly less total IR protein when compared to HFD (**Supplemental Fig. 1**). These initial findings suggested that the beneficial effects FEN treatment in LDLR-/- mice may have been more attributed to action in the liver than other insulin sensitive tissues.

### Fenretinide inhibits hepatic triglyceride accumulation and development of steatosis and alters hepatic metabolic gene expression in LDLR^-/-^ mice fed an atherogenic diet

LDLR^-/-^ mice are a recognised model of NAFLD, with high-fat feeding known to increase hepatic triglyceride accumulation (26). HFD resulted in a 2.5-fold increase in triglyceride content in the livers of LDLR^-/-^ mice (**Fig. 2A**). FEN treatment completely prevented intrahepatic triglyceride accumulation to levels similar to those in control mice. HFD feeding also led to a profound change in hepatic morphology compared to control mice including substantial lipid droplet accumulation (**Fig. 2B**). Whereas, FEN-HFD mice exhibited normal liver histology with the absence of lipid droplet accumulation within hepatocytes. Hepatic lipid homeostasis is maintained via a network of key transcription factors such as PPAIRα, LXR and SREBP which regulate the expression of genes involved in fatty acid synthesis, oxidation and transport. Dysregulation of this network in response to excess nutrition or genetic perturbations causes excess hepatic lipid accumulation and thus NALFD.

**Figure 2:**
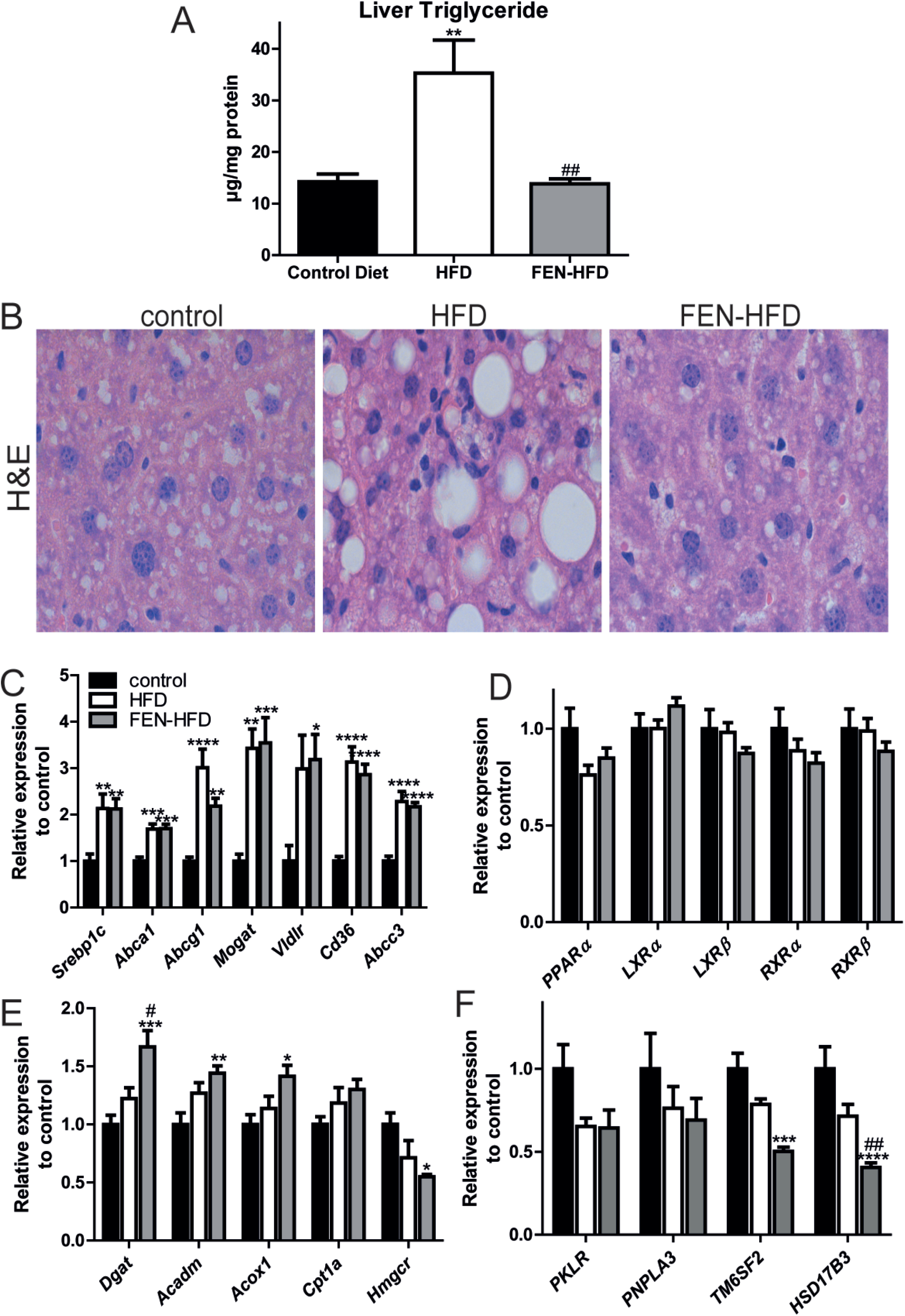
Fenretinide inhibits hepatic triglyceride accumulation and development of steatosis. (**A**) Hepatic triglyceride levels in LDLR^-/-^ mice. (**B**) Representative H&E staining of hepatic tissues. (**C-F)** Gene expression in liver tissue (n=8 per group). Data are represented as mean + S.E.M. and analysed by one-way ANOVA followed by Bonferroni multiple comparison t-tests where *p≤0.05, **p≤0.01, ***p≤0.001 and ****p≤0.0001 (control compared to HFD) or #p≤0.05 and ## p≤0.01 (HFD compared to FEN-HFD).

HFD (including high cholesterol) feeding +/− FEN of LDLR^-/-^ mice increased hepatic expression of LXR target genes, *Srebp1c, Abca1* and *Abcg1* and PPAIRα target genes *Mogat, Vldr, Cd36* and *Abcc3 (Mrp3)* compared to control diet mice (**Fig. 2C**). HFD +/− FEN did not affect the expression of *PPARα, LXR* or *RXR* transcription factors in liver (**Fig. 2D**). Hepatic *Dgat1, Acadm* and *Acox1*, also all PPAIRα target genes, were unaffected by HFD but were modestly increased by FEN treatment. *Cpt1* was not altered by either diet. However, FEN suppressed the statin target *Hmgcr* in liver (**Fig. 2E**) without affecting serum cholesterol levels (Table 1). HFD increased *Hmgcr* and *Abcc3* in white adipose tissue and FEN trended to prevent these increases, in addition, FEN suppressed adipose *Cd36* suggesting FEN also decreased adipose cholesterol in association with decreased adiposity (**Supplemental Fig. 1**).

Several human SNP/GWAS studies and more recent multi-omics data analyses have identified genes that have been described as key drivers of NAFLD (28). Of these, HFD did not affect the gene expression of hepatic *Pklr, Pnpla3, Tm6sf2* or *Hsd17b13* compared to control LDLR^-/-^ mice. However, FEN treatment resulted in a significant decrease in both *Tm6sf2* and *Hsd17b13* expression when compared to control mice (Figure 2F). FEN had no effect on *Pklr* or *Pnpla3* in this disease model (**Fig. 2F** *see discussion*).

### Fenretinide alters hepatic inflammatory and fibrotic gene expression

Persistent excess lipid accumulation is associated with a pro-inflammatory environment and the activation of hepatic stellate cells, the development of fibrosis and the progression to NASH, a more severe disease state. Indeed, HFD resulted in an increase in expression of the pro-inflammatory cytokine *TNFα*, the macrophage marker *Cd68* and profibrogenic signalling factor *TGF-β* that participates in hepatic stellate cell activation (29,30). FEN treatment significantly inhibited the increase in *Cd68* and trended to inhibit *TNFα* and *TGF-β* thus suggestive of a less pro-inflammatory, pro-fibrogenic environment (**Fig. 3A**). HFD did not alter the expression of *IL-6* or *Mcp-1* compared to control LDLR^-/-^ however, FEN treatment resulted in approximately 2.5-fold and 8-fold increase in gene expression respectively (**Fig. 3A**). HFD+/−FEN did not alter expression of anti-inflammatory gene, *IL-1*0.

**Figure 3:**
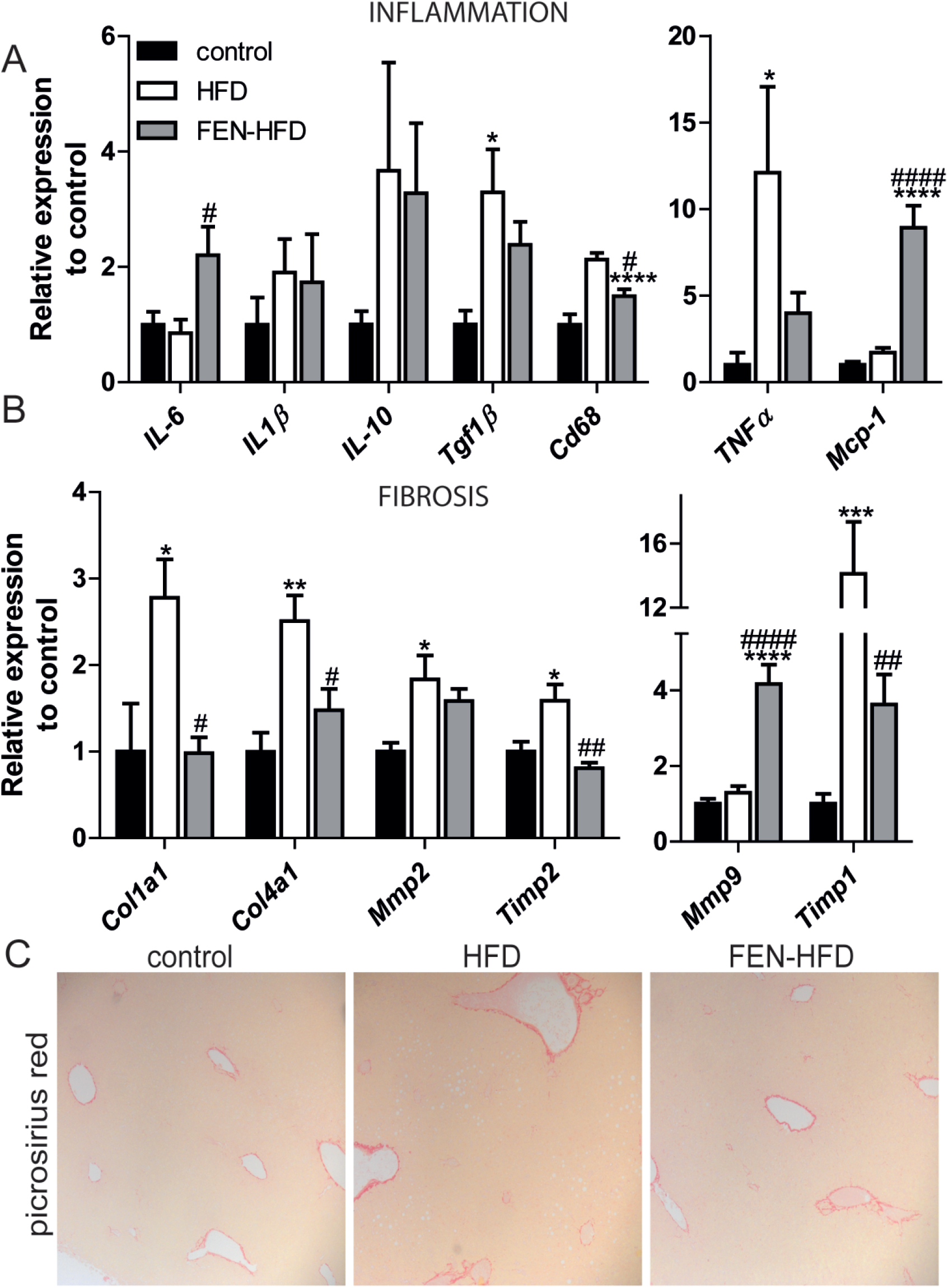
Fenretinide alters pro-inflammatory and fibrotic gene expression in liver. Hepatic mRNA expression of (**A**) inflammation and (**B**) fibrosis genes in LDLR^-/-^ mice (n=8 per group). (**C**) Representative picrosirius red staining of hepatic tissue. Data are represented as mean + S.E.M. and analysed by one-way ANOVA followed by Bonferroni multiple comparison t-tests where *p≤0.05, **p≤0.01, ***p≤0.001 and ****p≤0.0001 (control compared to HFD) or #p≤0.05 and ## p≤0.01 and #### p≤0.0001 (HFD compared to FEN-HFD).

HFD resulted in significant increases in the expression of genes driving fibrosis and tissue remodelling such as collagen (*Col1a1, Col4a1*), matrix metalloproteinases (*Mmp2*) and the tissue inhibitors of *Mmps (Timp1, Timp2;* **Fig. 3B**). FEN treatment almost completely inhibited the expression of all these genes to levels similar to those in control LDLR^-/-^ mice. A similar decrease in pro-inflammatory, macrophage and fibrosis genes with FEN treatment was determined in ApoE^-/-^ mice treated with HFD+/− FEN (**Supplemental Fig. 2**). In contrast to the inhibition of these genes (*TNFα, Col1a1etc*), FEN induced a 5-fold increase in *Mmp9* (Figure 3B) which is indicative of retinoid-specific signalling and increased clearance of pro-fibrotic proteins (31). However, despite these improvements in response to FEN treatment, there were no differences in hepatic fibrosis between all three diet groups in LDLR^-/-^ mice when determined by histologic stain picrosirius red (**Fig. 3C**).

### Fenretinide treatment inhibits *de novo* ceramide synthesis and lipotoxicity

Enzymes involved in ceramide biosynthesis, dihydroceramide desaturase, (DES1) and ceramide synthase (CerS)-6 have been implicated with increased ceramide production mediating obesity associated metabolic dysregulation in mice and humans (32). Hence, we next examined whether FEN treatment could inhibit the enzymes controlling *de novo* ceramide synthesis and thus lipotoxicity in the development of NAFLD/NASH in LDLR^-/-^ mice.

HFD increased DES 1 in LDLR^-/-^ mice (**Fig. 4A and 4B**), whereas FEN treatment prevented this increase so that protein levels were comparable to those control mice. Gene expression of hepatic dihydroceramide desaturase, *Degs1*, was unchanged with diet (**Fig. 4C**). However, HFD did trend to increase the hepatic *Cers6* and *Cers2* and FEN significantly decreased expression of *Cers6* (**Fig. 4C**).

**Figure 4:**
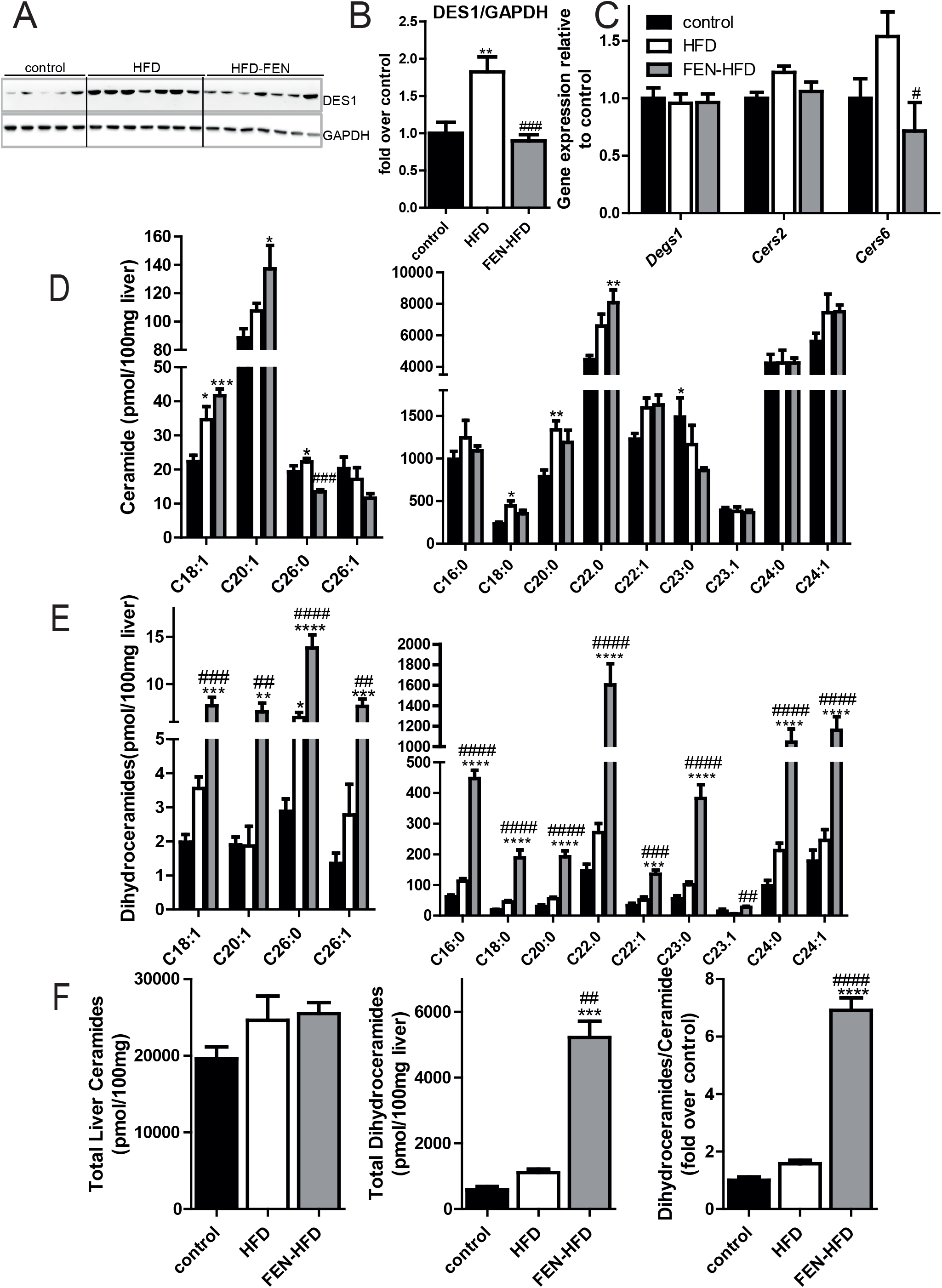
Fenretinide inhibits *de novo* ceramide synthesis via DES1 protein. (**A**) Western blot of liver tissue for DES1 (*upper panel*) and GAPDH (*lower panel*) used as a loading control, in LDLR^-/-^ mice. (**B**) Quantification of data shown in **A** (control n=10, HFD n=14, HFD-FEN n=14). (**C**) Hepatic mRNA expression of ceramide synthesis genes (n=8 per group). Quantification of ceramide (**D**), dihydroceramide (**E**) species in liver. (**F**) Total ceramide, dihydroceramide and ratio of total dihydroceramide: total ceramide in hepatic tissues (*left, middle and right panels respectively).* Data are represented as mean + S.E.M. and analysed by one-way ANOVA (**D, E, F**) followed by Bonferroni multiple comparison t-tests where *p≤0.05, **p≤0.01, ***p≤0.001 and ****p≤0.0001 (control compared to HFD) or #p≤0.05 and ## p≤0.01, ### p≤0.001 and #### p≤0.001 (HFD compared to FEN-HFD).

HFD increased several acyl ceramide species e.g., C18:0, C18:1 and C20:0 but total ceramide levels were not increased with HFD+/−FEN compared to control LDLR^-/-^. FEN treatment specifically decreased C26:0 ceramide but not any other species (**Fig. 4D**). However, FEN treatment increased all species of dihydroceramides measured from C16:0 to C26:1 and total dihydroceramide levels by 4.7 to 8.9-fold compared to HFD and control mice respectively. Similar results were obtained in male and female ApoE^-/-^ mice (**Supplemental Fig. 3**).

HFD also elevated levels of the ER stress protein GRP78/BIP (**Supplemental Fig. 4**) and FEN almost completely inhibited this increase. eIF2α phosphorylation and CHOP protein expression trended to be altered similarly, but HFD+/−FEN did not affect levels of autophagy proteins beclin-1 and p38 (**Supplemental Fig. 4**). Taken together, these results suggest that FEN mediated inhibition of DES1 protein and thus inhibition of excess ceramide biosynthesis and lipotoxicity may be part of the mechanism of preventing insulin resistance and NAFLD/NASH.

### Fenretinide worsens hypertriglyceridemia and accelerates atherogenesis in LDLR^-/-^ mice

The liver packages triglycerides into lipoproteins together with cholesterol and apolipoproteins which are then transported in the circulation. Thus, next we investigated this system to determine the effects of HFD+/−FEN on development of dyslipidemia and atherosclerosis. HFD caused a major increase in circulating triglycerides and total cholesterol compared to control LDLR^-/-^ mice (**Fig. 5A, 5B**). Surprisingly, FEN did not prevent the increase in serum cholesterol and caused a further increase in serum triglyceride when compared to HFD mice. HFD elevated circulating apolipoprotein B (ApoB) 48 levels, but FEN-HFD resulted in increased ApoB100 protein in both, liver and serum (**Fig. 5C-5E**) suggestive of unique effects respectively on further increasing very low-density lipoprotein (VLDL) and/or LDL levels in LDLR^-/-^ mice.

**Figure 5:**
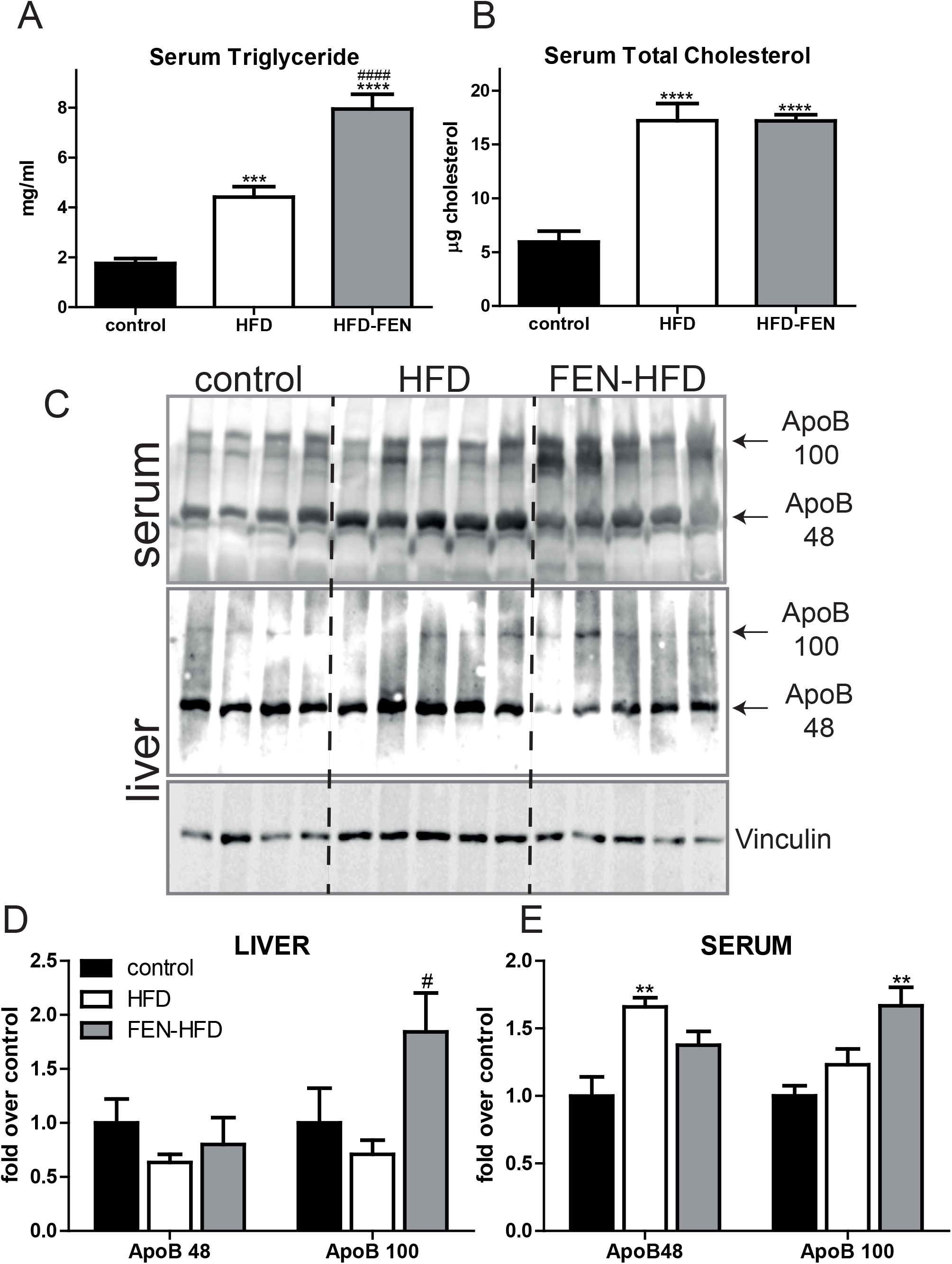
Fenretinide increases circulating triglyceride and apolipoprotein B 100 levels. Serum triglyceride (**A**) and total cholesterol (**B**) in LDLR^-/-^ mice (n=8 per group). (**C**) Western blot of ApoB 48 and ApoB 100 in serum (upper panel) and hepatic tissues (middle panel). Quantification of hepatic (**D**) and serum levels (**E**) shown in (**C**). Hepatic levels were normalised to vinculin. Data are represented as mean + S.E.M. (n=4-5 per group) and analysed by one-way ANOVA followed by Bonferroni multiple comparison t-tests where **p≤0.01, ***p≤0.001 and ****p≤0.0001 (control compared to HFD) or #p≤0.05 and ####p≤0.0001 (HFD compared to FEN-HFD).

Since elevated circulating triglyceride, ApoB-containing lipoproteins and the ratio of ApoB100 to ApoB48 are major risk factors for the development of CVD (33), we next investigated the effect of HFD+/−FEN on atherosclerotic plaque formation. HFD resulted in atherosclerotic plaque formation in the aortic root, in the aortic arch and the descending aorta in LDLR^-/-^ mice (**Fig. 6A and 6B**). FEN-treated mice had a similar level of plaque formation compared to HFD mice in the aortic root and in the aortic arch, but considerably more atherosclerotic plaque throughout the descending aorta (**Fig. 6A and 6B**). To determine if this was the case in another commonly used model for atherosclerosis, we examined plaque formation in ApoE^-/-^ mice and found it to be accelerated in this background too. FEN-HFD resulted in significantly greater plaque accumulation in the descending aorta of female mice (**Supplemental Fig. 5**).

**Figure 6:**
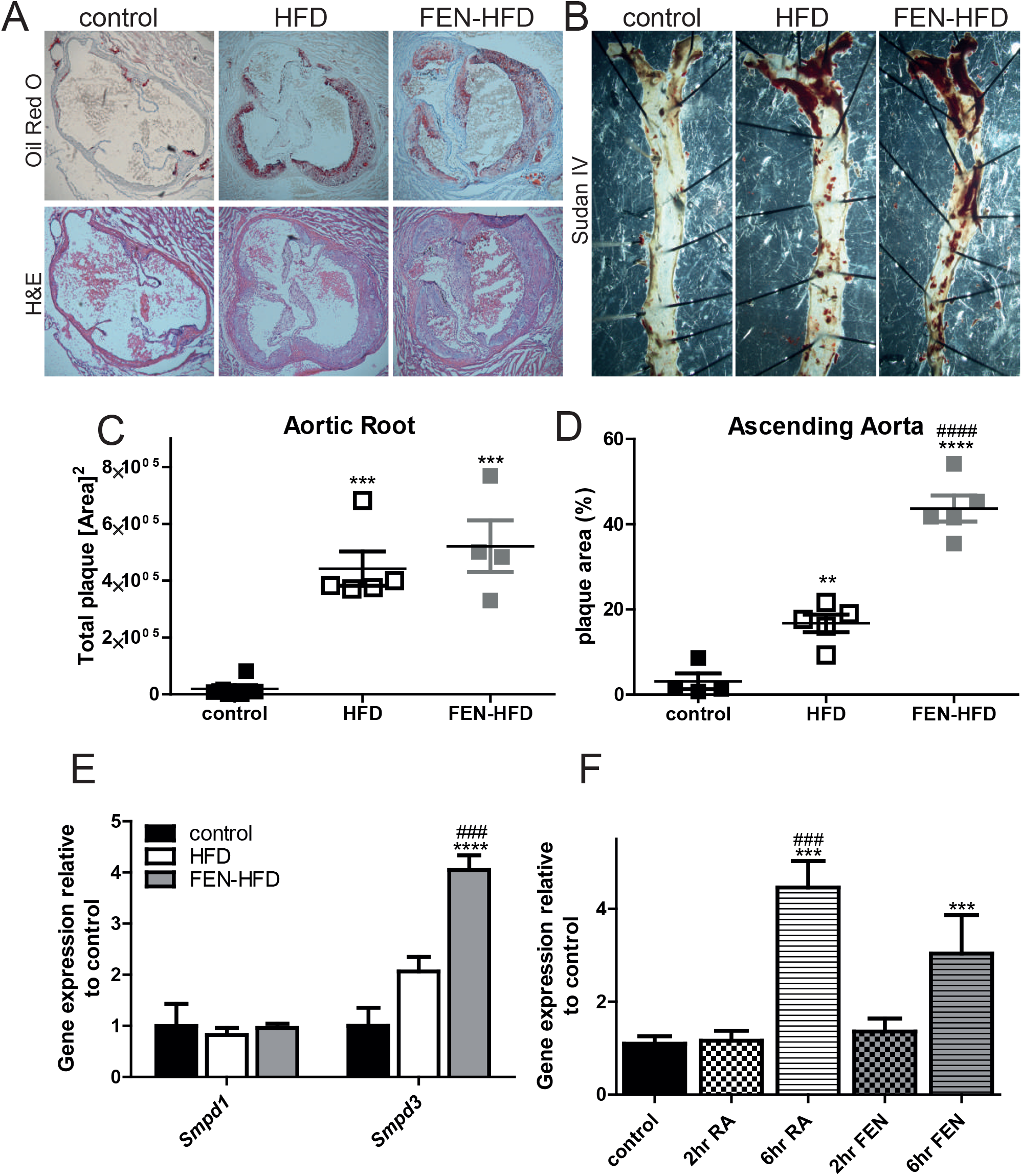
Fenretinide accelerates atherogenesis in LDLR^-/-^mice. (**A**) Plaque formation (upper panels) in heart aortic roots sections stained with Oil Red O and overall structure with H&E staining (lower panels). (**B**) Plaque formation in the decending aorta prepared *en face* and stained with Sudan IV. (**C**-**D**) Quantification is shown in A and B respectively. (**E)** Hepatic expression of genes encoding neutral sphingomyelinases, *Smpd1* and *Smpd3* and WAT expression of *Smpd3* (n=8 per group). (**F**) Hepatic *Smpd3* gene expression following acute retinoic acid (RA) or FEN intraperitoneally injection in C57BL/6 mice (n=8 per group). Data are represented as mean + S.E.M. and analysed by One-way ANOVA followed by Bonferroni multiple comparison t-tests where **p≤0.01, ***p≤0.001 and ****p≤0.0001 (compared to control) or ### p≤0.001 and #### p≤0.0001 (HFD compared to HFD-FEN).

In addition to the role of excess *de novo* ceramide synthesis in the pathogenesis of metabolic diseases, ceramide generation via sphingomyelin hydrolysis has also been linked to atherosclerosis (see Choi *et al* for recent review (10)). FEN treatment in LDLR^-/-^ mice lead to a striking 4-fold increase in hepatic *Smpd3* expression, the gene encoding neutral Sphingomyelinase-2, recently shown to contribute to the development of atherosclerosis in ApoE^-/-^ mice. (**Fig. 6E**). Similar results were obtained in our ApoE^-/-^ mice (**Supplemental Fig. 5**). *Smpd3* expression was not altered in white adipose tissue (**Fig. 6E**). We tested whether *Smpd3* could be induced by FEN directly via RAR-signalling, to understand the mechanism behind this alteration. Acute RA injection led to a potent increase in *Smpd3* expression at 6 hours but not earlier at 2 hours in the livers of lean C57/BL6 mice. FEN treatment also led to an increase in *Smpd3* expression at 6 hours, but the effect was not as striking as with RA treatment (**Fig. 6F**).

Next, we examined whether an increase in hepatic *Smpd3* expression could result in an increase in circulating ceramides and thereby contribute to the increased development of atherosclerosis FEN treatment in LDLR-/- mice. FEN increased total serum ceramide levels 1.6-fold more than in HFD mice (Figure 7). We determined an increase in a number of ceramide species from FA acyl groups 18:0 to 26:0. FEN also increased total serum dihydroceramide levels 8-fold higher than in HFD mice with increases in every species measured (Fig. 7).

**Figure 7:**
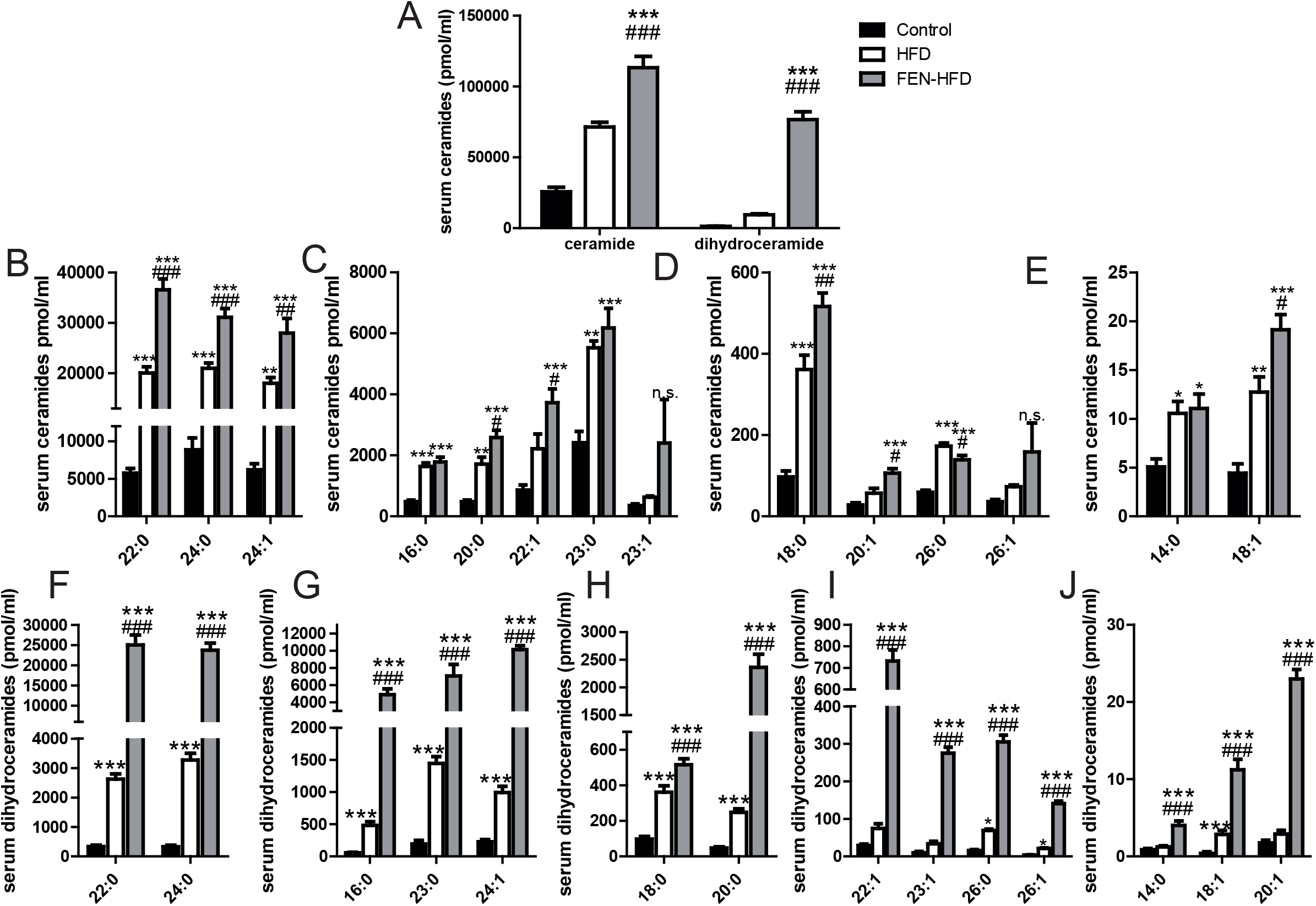
Fenretinide increases serum ceramide and dihydroceramide levels in LDLR^-/-^ mice. Quantification of ceramide (**A, B-E**) and dihydroceramide (**A, F-J**) species in serum. Data are represented as mean + S.E.M. and analysed by one-way ANOVA followed by Bonferroni multiple comparison t-tests where *p≤0.05, **p≤0.01 and ***p≤0.001 (control compared to HFD) or #p≤0.05, ## p≤0.01 and ### p≤0.001 (HFD compared to FEN-HFD).

Thus, overall, these data suggest that FEN treatment was beneficial in the treatment of pathologies associated with an obesogenic diet and excess fat gain, thereby attenuating the development of insulin resistance lipotoxicity and NAFLD/NASH, but at the detriment of the cardiovascular system, at least in genetic mouse models lacking LDLR^-/-^ or ApoE^-/-^. Mechanistically, FEN treatment results in retinoic acid signalling mediated induction of sphingomyelinase gene *Smpd3* and an increase in circulating ceramides and thereby contributes to the increased development of atherosclerosis in LDLR-/- mice (as illustrated in Fig. 8).

**Figure 8:**
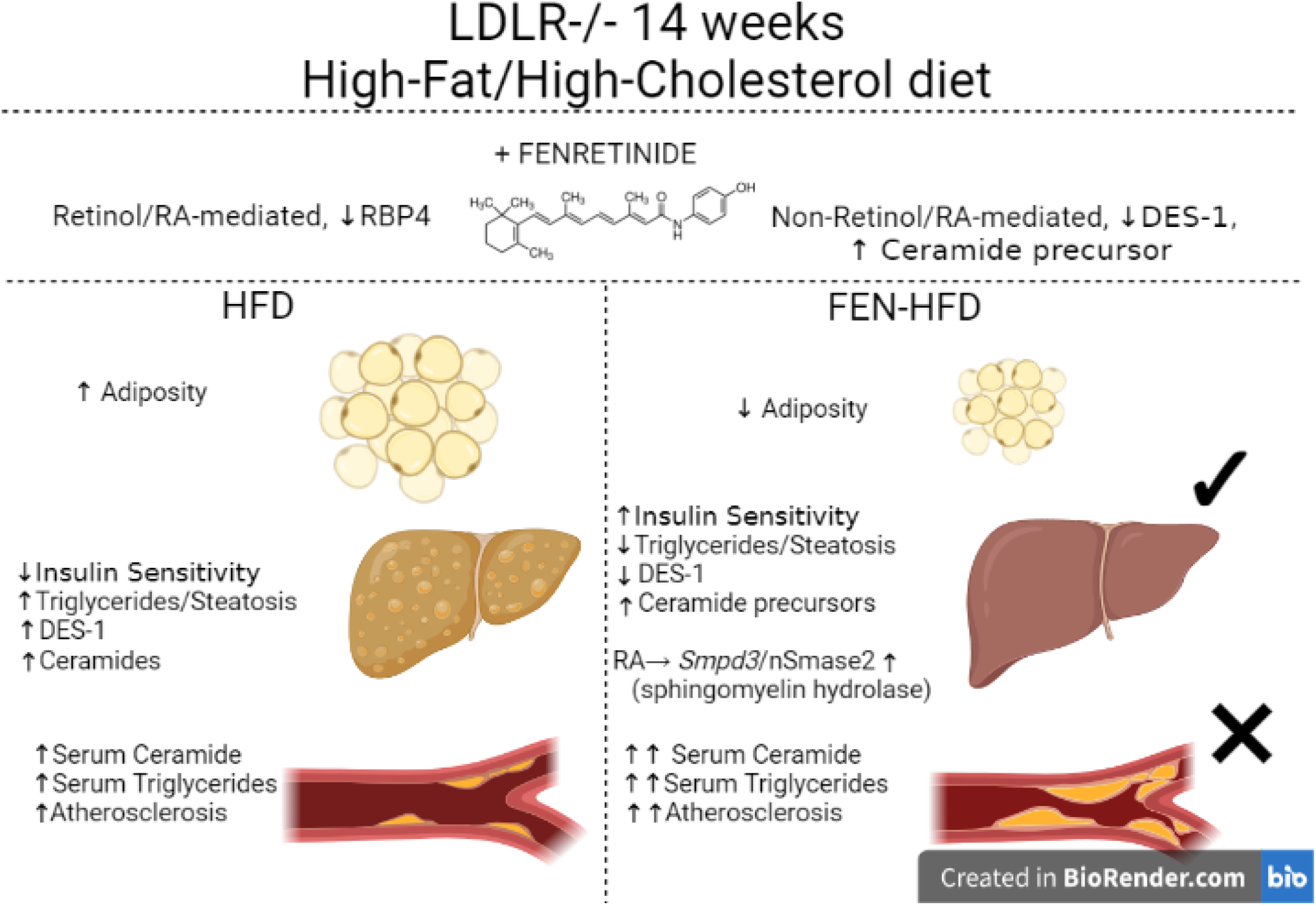
Working Model of Fenretinide Action. A graphical representation demonstrating the effects of FEN on multiple metabolic complications using LDLR^-/-^ mice fed high-fat/high-cholesterol diet +/− FEN as a model of atherosclerosis and non-alcoholic fatty liver disease (NAFLD). FEN treatment prevented excess weight gain and adiposity, improved insulin sensitivity and completely inhibited hepatic triglyceride accumulation, ballooning and steatosis. In association, FEN decreased the expression of hepatic genes driving NAFLD, inflammation and fibrosis e.g., *Hsd17b13, Cd68* and *Col1a1.* The mechanisms of FEN’s beneficial effects in association with decreased adiposity were mediated by suppression of ceramide synthesis, via hepatic DES1 protein, leading to increased ceramide precursors (i.e. dihydroceramides). However, FEN treatment in LDLR^-/-^ mice enhanced circulating triglycerides and accelerated aortic plaque formation. FEN increased hepatic sphingomyelinase *Smpd3* expression, via a retinoic acid mediated mechanism, linking induction of an alternative ceramide generation pathway (via sphingomyelin hydrolysis) to a novel mechanism of increased atherosclerosis. Pharmacological treatment of both DES1 and *Smpd3/nSMase2* may be a novel, more potent therapeutic approach for the treatment of metabolic syndrome.

## DISCUSSION

The multiple overlapping secondary pathologies in response to diet-induced obesity such as the development of metabolic syndrome, type 2 diabetes, NAFLD, atherosclerosis and CVD have interconnected underlying molecular drivers (1,2,5,34). There is increasing evidence that dysregulation of lipid metabolism e.g., excess ceramide biosynthesis that can lead to lipotoxicity and insulin resistance, may be a tractable target for novel pharmacological interventions that can treat these co-morbidities. Currently there is no treatment for NAFLD or NASH and therapies are urgently needed. In this study we have demonstrated that inhibition of adiposity and inhibition of DES1, the final step in ceramide biosynthesis, by FEN can prevent hepatic triglyceride accumulation and steatosis in the LDLR^-/-^ mouse model of diet-induced NAFLD and atherosclerosis. This comes at the expense of the cardiovascular system however, as FEN treatment results in augmented atherogenesis accompanied by alterations in hepatic and serum ApoB 100 species of lipoproteins and induction of *Smpd3* (as illustrated in Fig. 8).

It has previously been demonstrated that genetic deletion of enzymes involved in ceramide production and pharmacological treatment with myriocin, a natural fungal metabolite that inhibits serine palmitoyltransferase can attenuate obesity and improve glucose homeostasis (11,35). Many of these studies have also shown an associated attenuation of hepatic steatosis (13,15). We have previously demonstrated that FEN can partially prevent diet-induced obesity and associated metabolic pathologies, including a partial attenuation of fatty liver (15,16). In genetically obese (leptin-signalling deficient) mice, FEN markedly improved glucose homeostasis without a decrease in body weight or liver triglyceride (17). Thus, it is even more striking that FEN can prevent hepatic triglyceride accumulation and steatosis in LDLR^-/-^ mice, a more robust model of dyslipidemia and NAFLD compared to obesity-prone C57BL/6 mice (26). FEN did not improve glucose homeostasis in LDLR^-/-^ mice, despite inhibition of ceramide biosynthesis and improvement in insulin action. Thus together, these findings suggest overlapping pathologies of metabolic syndrome can be dissociated depending on the animal model investigated and this can affect the beneficial effects observed with therapeutic treatments such as FEN.

Hepatic gene expression alterations that are linked with excess hepatic lipid accumulation revealed a unique pattern of beneficial improvement with FEN treatment including downregulation of *Tm6sf2* and *Hsd17b13* but not *Mogat* or *Vldr.* The biological roles of *Tm6sf2* and *Hsd17b13* appear to involve apolipoprotein secretion and lipid droplet homeostasis, respectively, but these are not clear and require further investigation to understand their role in normal lipid metabolism and dyslipidemia (3). FEN treatment also decreased genes involved in deposition of excess extracellular matrix such as *Col1a1* and *Timp2*, but increased *Mmp9* expression (31). *Mmp9* has been reported to be a directly retinoid-responsive gene and confirmed by us to be directly regulated by RA and FEN treatment by RNA-seq and qPCR methods (*not shown, manuscript in preparation).* This induction of *Mmp9* may directly contribute to a retinoid-specific effect to increase degradation of extracellular matrix and thus prevent the progression of hepatic fibrosis (31).

FEN markedly increased aortic atherosclerotic plaque formation in LDLR^-/-^ mice, despite all the beneficial effects of FEN and the demonstration that myriocin-mediated inhibition of sphingolipid biosynthesis decreased atherosclerosis in ApoE^-/-^ mice (36). In those studies, myriocin also decreased plasma triglycerides in hyperlipidemic ApoE^-/-^ mice, whereas here FEN elevated circulating triglyceride, ApoB-containing lipoproteins and the ratio of ApoB100 to ApoB48. Since these are major risk factors for the development of atherosclerosis, this may contribute to the mechanism of increased aortic plaque formation. Some retinoic acid derivatives (that act primarily via retinoic X receptor, RXRs), have been used as an acne medication and in some patients results in hypertriglyceridemia via regulation of lipoproteins and thus careful monitoring is required (37–39). Here, FEN treatment increased levels of ApoB 100 in both, the liver and serum, but not hepatic gene expression of other apolipoprotein species and moreover FEN and RA act via RARs not RXRs. FEN may regulate apolipoprotein B secretion via downregulation of TM6SF2, which was recently identified as a ChREBP target in mouse liver but there is no evidence that FEN or other synthetic RA derivatives can regulate ChREBP activity (17) and manuscript under preparation).

Our data suggest the mechanism of increased atherosclerosis with FEN treatment is via the upregulation of the gene encoding type 2-neutral sphingomyelinase (nSMase 2, gene *Smpd3*), an alternative ceramide generation pathway (via sphingomyelin hydrolysis). Genetic deficiency of nSMase2 (in mutant *Smpd3*^fro/fro^ mice) or pharmacological inhibition of nSMase2 activity significantly reduced the size of atherosclerotic lesions in ApoE^-/-^ mice (40,23). nSMase2 is a key enzyme of sphingolipid metabolism and *Smpd3* expression is highest in the brain but also significant in the liver (40). Interestingly, *Smpd3* has been identified as a RA induced gene in MCF7 breast carcinoma cells and mouse embryonic stem cells treated for 12-24 h with retinoic acid and nSMase2 activity is regulated a number of factors eg. pro-inflammatory cytokines and phosphorylation (21,22,41). More recently, Jiang & co-workers (42) reported that upregulation of intestinal *Smpd3* induces intestinal ceramide production and secretion to increase circulating ceramide levels resulting in accelerated atherosclerosis. The upregulation of intestinal *Smpd3* was attributed to farnesoid X receptor (FXR), a ligand-activated nuclear receptor that regulates cholesterol and bile acid metabolism. Both studies reported no influence on serum cholesterol or triglyceride levels (23,42)

Increased aortic atherosclerosis with FEN treatment was recapitulated in the ApoE^-/-^ mice (Supplemental material) and most recently by Chiesa and co-workers (43), but in contrast they reported a decrease in circulating triglyceride and lipoprotein levels. In humans, FEN treatment over two years also prevented an increase in circulating triglyceride levels (associated with an increase in HOMA) in normal weight women (44). It is currently unclear whether FEN, either via retinoid signalling or inhibition of ceramide biosynthesis can cause hypertriglyceridemia. The effect of DES1 genetic knockout has not been studied in ApoE^-/-^ or LDLR^-/-^ and may help to clarify this. Chiesa and co-workers study of FEN in ApoE^-/-^ mice also reported splenomegaly and haematological alterations (43). In our study, although FEN treatment also resulted in splenomegaly in both male and female ApoE^-/-^ mice, it did not in LDLR^-/-^ mice (**Supplemental Fig. 6**). Thus, we do not attribute increased atherosclerotic lesions to haematological defects.

In summary, the present study has demonstrated that FEN treatment can prevent hepatic triglyceride accumulation, steatosis and fibrosis in addition to prevention of obesity in LDLR^-/-^ mice. Part of this favourable effect is via prevention of obesity and also inhibition of ceramide biosynthesis and improvement in insulin action. Despite these beneficial metabolic effects, a clear worsening of atherosclerosis was established in this atherosclerosis prone mouse model, which appears to involve an alternative ceramide generation pathway (via sphingomyelin hydrolysis) regulated by *Smpd3*/nSMase2. Since excess ceramide production causes lipotoxicity, metabolic dysregulation and atherogenesis, dual targeting of both DES1 and *Smpd3*/nSMase2 may be a novel strategy for treatment of metabolic syndrome and importantly deadly co-morbidities.

## Acknowledgements

This study was supported by funds from the British Heart Foundation (PG16/90/32518) project grant to N. Mody and a PhD studentship to S.M. by the James Mearns Trust and School of Medicine, Medical Sciences and Nutrition, University of Aberdeen (UoA).

No potential conflicts of interest relevant to this article are reported.

N. Mody and D.T. conceived the study, designed the experiments and wrote the manuscript. D.T.,

S.M, N. Morrice, R.D., S.K.S, E.F. N.A., P.H. performed experiments. P.W. and M.D. performed the ceramide analysis, M.D. contributed to the study conception, discussions and reviewed the manuscript.

N.M. is the guarantor of this work and, as such, had full access to all the data in this study and takes responsibility for the integrity and accuracy of the data analysis.

The authors thank the UoA animal research facility, qPCR core facility (Institute of Medical Sciences, UoA) and Linda Davidson (NHS Grampian) for their technical contributions regarding animal studies, qPCR and histology respectively and Alison Rutter at UHI for lipidomics support. The authors also wish to thank Patrick W F Hadoke (University of Edinburgh) and Heather M Wilson (UoA) for invaluable discussions about mechanisms of atherosclerosis and review of the manuscript, Matteo Zanda, Sergio Dall’Angelo, Chiara Zanato and Ilaria Patruno (all UoA) for medicinal chemistry support.

