## Supplemental Material, Table 1 and Figures 1-6 for "Fenretinide inhibits obesity and fatty liver disease but induces Smpd3 to increase serum ceramides and worsen atherosclerosis in LDLR-/- mice"

**Supplemental Material and Methods**

**Immunoblotting.** Tissues were prepared as previously described in materials and methods. Membranes were probed for the following additional proteins; phospho-p38 (The 181/Tyr 182, cat: 4581), total p-38 (cat: 8690), phospho-eif2α (Ser 51, cat: 3398), total eif2α (cat: 5321), beclin 1 (cat: 3495) and GAPDH (cat: 5174, all Cell Signaling Technology), or IR β-chain, Bip and CHOP (Santa Cruz Biotechnology, cat: sc-373975, sc166490 and sc-7531 respectively).

### Supplemental Figures

**Supplemental Table 1:** qPCR primer details.

| Gene | Forward primer (5’-3’) | Reverse Primer (5’-3’) |
|---|---|---|
| ***Abca1*** | AAAACCGCAGACATCCTTCAG | CATACCGAAACTCGTTCACCC |
| ***Abcc3*** | CTGGGTCCCCTGCATCTAC | GCCGTCTTGAGCCTGGATAAC |
| ***Abcg1*** | GTGGATGAGGTTGAGACAGACC | CCTCGGGTACAGAGTAGGAAAG |
| ***Acadm*** | AGGGTTTAGTTTTGAGTTGACGG | CCCCGCTTTTGTCATATTCCG |
| ***Acox1*** | TAACTTCCTCACTCGAAGCCA | AGTTCCATGACCCATCTCTGTC |
| ***Adiponectin*** | TGTTCCTCTTAATCCTGCCCA | CCAACCTGCACAAGTTCCCTT |
| ***ApoA1*** | GGCACGTATGGCAGCAAGAT | CCAAGGAGGAGGATTCAAACTG |
| ***ApoA2*** | GCAGACGGACCGGATATGC | GCTGCTCGTGTGTCTTCTCA |
| ***ApoC3*** | TACAGGGCTACATGGAACAAGC | CAGGGATCTGAAGTGATTGTCC |
| ***ApoE*** | CTGACAGGATGCCTAGCCG | CGCAGGTAATCCCAGAAGC |
| ***Cd36*** | GAACCACTGCTTTCAAAAACTG G | TGCTGTTCTTTGCCACGTCA |
| ***Cd68*** | TGTCTGATCTTGCTAGGACCG | GAGAGTAACGGCCTTTTTGTGA |
| ***Cers2*** | AAGTGGGAAACGGAGTAGCG | ACAGGCAGCCATAGTCGTTC |
| ***Cers6*** | CGGCTGGGCATATTTCCTCT | GTCATCCTTGGATACCTTGCCT |
| ***Col1a1*** | CCAAAGGTGCTGATGGTTCT | ACCAGCTTCACCCTTGTCAC |
| ***Col4a1*** | GCTCTGGCTGTGGAAAATGT | CTTGCATCCCGGGAAATC |
| ***Cpt1a*** | CTCCGCCTGAGCCATGAAG | CACCAGTGATGATGCCATTCT |
| ***Cyp26A1*** | TTCGGGTTGCTCTGAAGACT | TCCTCCAAATGGAATGAAGC |
| ***Degs1*** | TCCCTACTCGCGGATGAAGA | TTTGAAGCCGTGGACAGGAA |
| ***Dgat*** | TCCGTCCAGGGTGGTAGTG | TGAACAAAGAATCTTGCAGACGA |
| ***G6Pase*** | ATGAACATTCTCCATGACTTTGGG | GACAGGGAACTGCTTTATTATAGG |
| ***GLUT4*** | GGAAGGAAAAGGGCTATGCTG | TGAGGAACCGTCCAAGAATGA |
| ***Hmgcr*** | GATTCTGGCAGTCAGTGGGAA | GTTGTAGCCGCCTATGCTCC |
| ***HSD17B13*** | ATTCCCCGGAGAAGGAAATCT | CAGCCTGCCTATTCCGTGT |
| ***IL-10*** | GCTCTTACTGACTGGCATGAG | CGCAGCTCTAGGAGCATGTG |
| ***IL1β*** | GCAACTGTTCCTGAACTCAACT | ATCTTTTGGGGTCCGTCAACT |
| ***IL-6*** | TAGTCCTTCCTACCCCAATTTCC | TTGGTCCTTAGCCACTCCTTC |
| ***LRAT*** | CCGTCCCTATGAAATCAGCTC | ATGGGCGACACGGTTTTCC |
| ***LXRα*** | CTCAATGCCTGATGTTTCTCCT | TCCAACCCTATCCCTAAAGCAA |
| ***LXRβ*** | GCCTGGGAATGGTTCTCCTC | AGATGACCACGATGTAGGCAG |
| ***Mcp-1*** | TTAAAAACCTGGATCGGAACCAA | GCATTAGCTTCAGATTTACGGGT |
| ***Mmp2*** | CAAGTTCCCCGGCCATGTC | TTCTTGGTCAAGCTCACCTGTC |
| ***Mmp9*** | GCGTCGTGATCCCCACTTAC | CAGGCCGAATAGGAGCGTC |
| ***Mogat1*** | TGGACGCCAGTTTGGTTCCAG | TGCTCTGAGGTCGGGTTCA |
| ***Nono*** | GCCAGAATGAAGGCTTGACTAT | TATCAGGGGGAAGATTGCCCA |
| ***PEPCK*** | GAGATAGCGGCACAAT | TTCAGAGACTATGCGGTG |
| ***PNPLA3*** | TCACCTTCGTGTGCAGTCTC | CCTGGAGCCCGTCTCTGAT |
| ***Pparα*** | ACGATGCTGTCCTCCTTGATG | GTGTGATAAAGCCATTGCCGT |
| ***Pparγ*** | AGTGGAGACCGCCCAGG | GCAGCAGGTTGTCTTGGATGT |
| ***RARα*** | CGC CAA GGG AGC TGA ACG GG | GGG TGG CTG GGC TGC TTC TG |
| ***RARβ*** | CGCGAGCCCTTCCTCCTGC | AAAAGCCCTTGCACCCCTCGC |
| ***Resistin*** | AAGAACCTTTCATTTCCCCTCCT | GTCCAGCAATTTAAGCCAATGTT |
| ***RPB4*** | ACGAGTCCGTCTTCTGAGCAACTG | GCACAGCTCCTCCTGCCGTT |
| ***RXRα*** | ATGGACACCAAACATTTCCTGC | CCAGTGGAGAGCCGATTCC |
| ***RXRβ*** | CCACCTCTTACCCCTTCAGC | TGGAAGAACTGATGACTGGGA |
| ***Smpd1*** | TGGGACTCCTTTGGATGGG | CGGCGCTATGGCACTGAAT |
| ***Smpd3*** | ACACGACCCCCTTTCCTAATA | GGCGCTTCTCATAGGTGGTG |
| ***Srebp1c*** | GATGTGCGAACTGGACACAG | CATAGGGGGGGTCAAACAG |
| ***Tgf1β*** | AGCCCGAAGCGGACTACTAT | CTGTGTGAGATGTCTTTGGTTTTC |
| ***Timp1*** | GCAACTCGGACCTGGTCATAA | CGGCCCGTGATGAGAAACT |
| ***Timp2*** | TCAGAGCCAAAGCAGTGAGC | GCCGTGTAGATAAACTCGATGTC |
| ***TM6SF2*** | AGTTTCGGCGTTCTCACAGC | GCATAGAGAGGGTCGTAGGTG |
| ***TNFα*** | CCCTCACACTCAGATCATCTTCT | GCTACGACGTGGGCTACAG |
| ***Vldlr*** | GGCAGCAGGCAATGCAATG | GGGCTCGTCACTCCAGTCT |
| ***Ywaz*** | GAAAAGTTCTTGATCCCCAATG C | TGTGACTGGTCCACAATTCCTT |

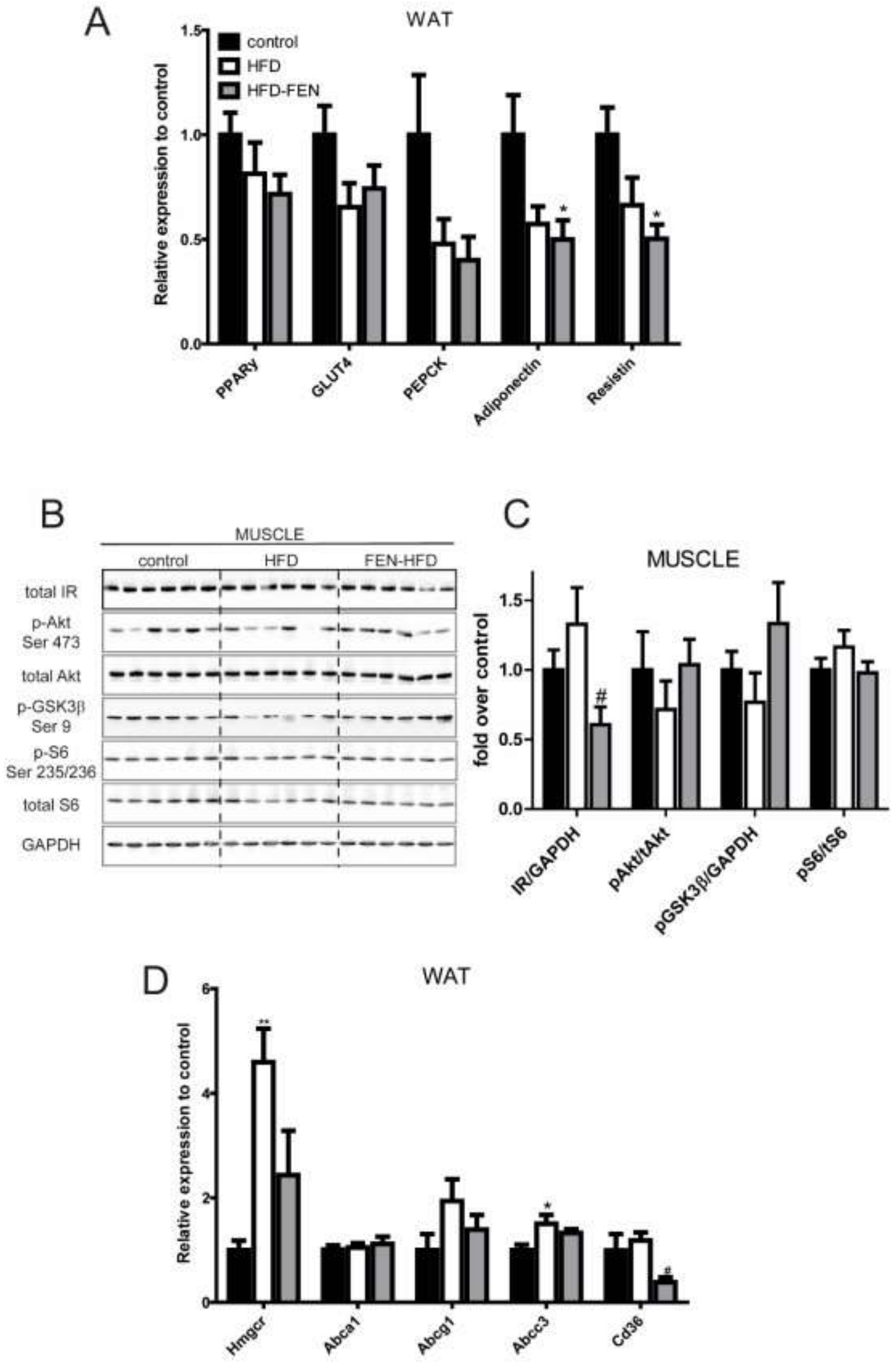


Supplemental Figure 1

**Supplemental Figure 1: WAT and skeletal muscle analysis in LDLR$^{-/-}$ mice.** Genetic analysis of (**A, D**) WAT in LDLR$^{-/-}$ mice (n=8 per group) as analysed by qPCR using SYBR green and LightCycler 480 (Roche). (B) Western blot of skeletal muscle from LDLR-/- mice and quantification shown in (**C**). Data are represented as mean + S.E.M. and analysed by one-way ANOVA followed by Bonferroni multiple comparison t-tests where *$p \leq 0.05$ (control compared to HFD) or #$p \leq 0.05$ (HFD compared to FEN-HFD).

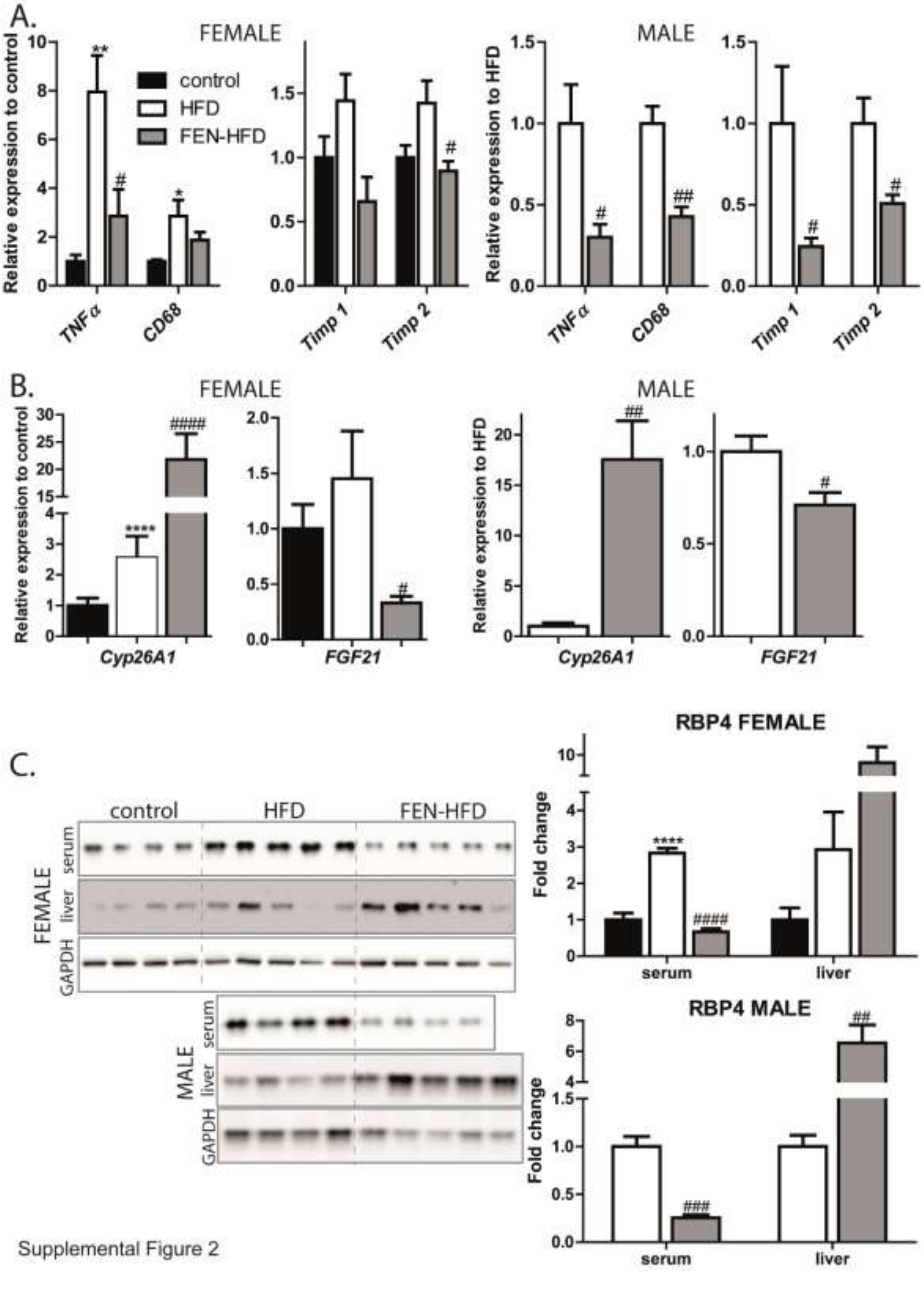


Supplemental Figure 2

**Supplemental Figure 2: Fenretinide alters pro-inflammatory and fibrotic genes and RBP4 levels in male and female ApoE-/- mice.** (**A, B**) Genetic analysis of hepatic tissues in male and female ApoE$^{-/-}$ mice (control n=4, HFD n=5, FEN-HFD n=5). (**C**) Western blot analysis of RPB4 levels in serum and hepatic tissues from female (upper left panels) and male (lower left panels) and quantification (right panels). Data are represented as mean + S.E.M. and analysed by one-way ANOVA followed by Bonferroni multiple comparison t-tests where *p≤0.05, **p≤0.01 and ****p≤0.0001 (compared to control) or #p≤0.05, ##p≤0.01, ### p≤0.001 and #### p≤0.0001 (HFD compared to HFD-FEN).

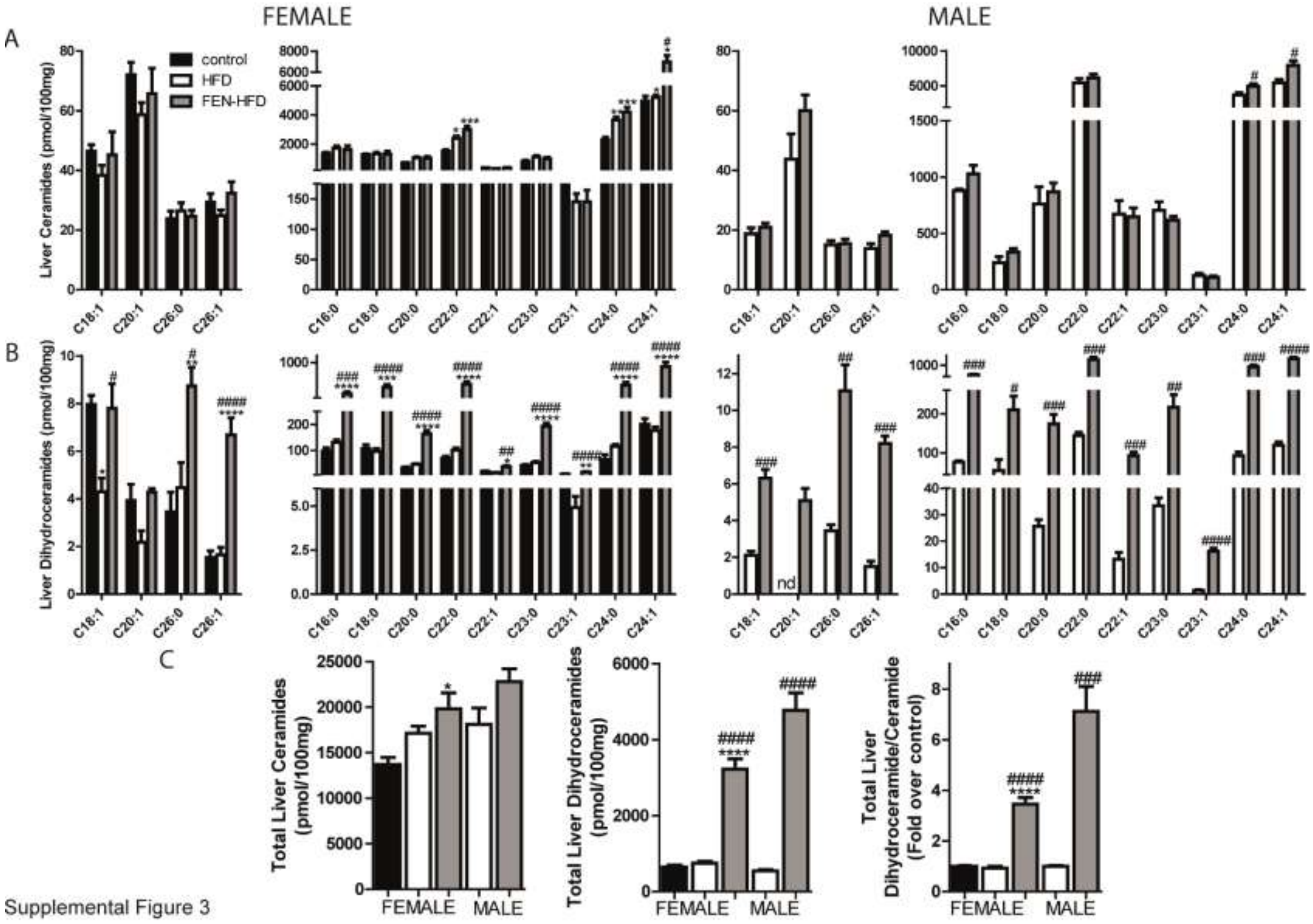


Supplemental Figure 3

**Supplemental Figure 3: Fenretinide increases dihydroceramide species in hepatic tissues of male and female ApoE$^{-/-}$ mice.** Lipidomic analysis of Ceramide (**A**), Dihydroceramide (**B**) and total (**C**) species in hepatic tissues. Data are represented as mean + S.E.M. and analysed by one-way ANOVA followed by Bonferroni multiple comparison t-tests where *p≤0.05, **p≤0.01, ***p≤0.001 and ****p≤0.0001 (control compared to HFD) or #p≤0.05 and ## p≤0.01, ### p≤0.001 and #### p≤0.001 (HFD compared to FEN-HFD).

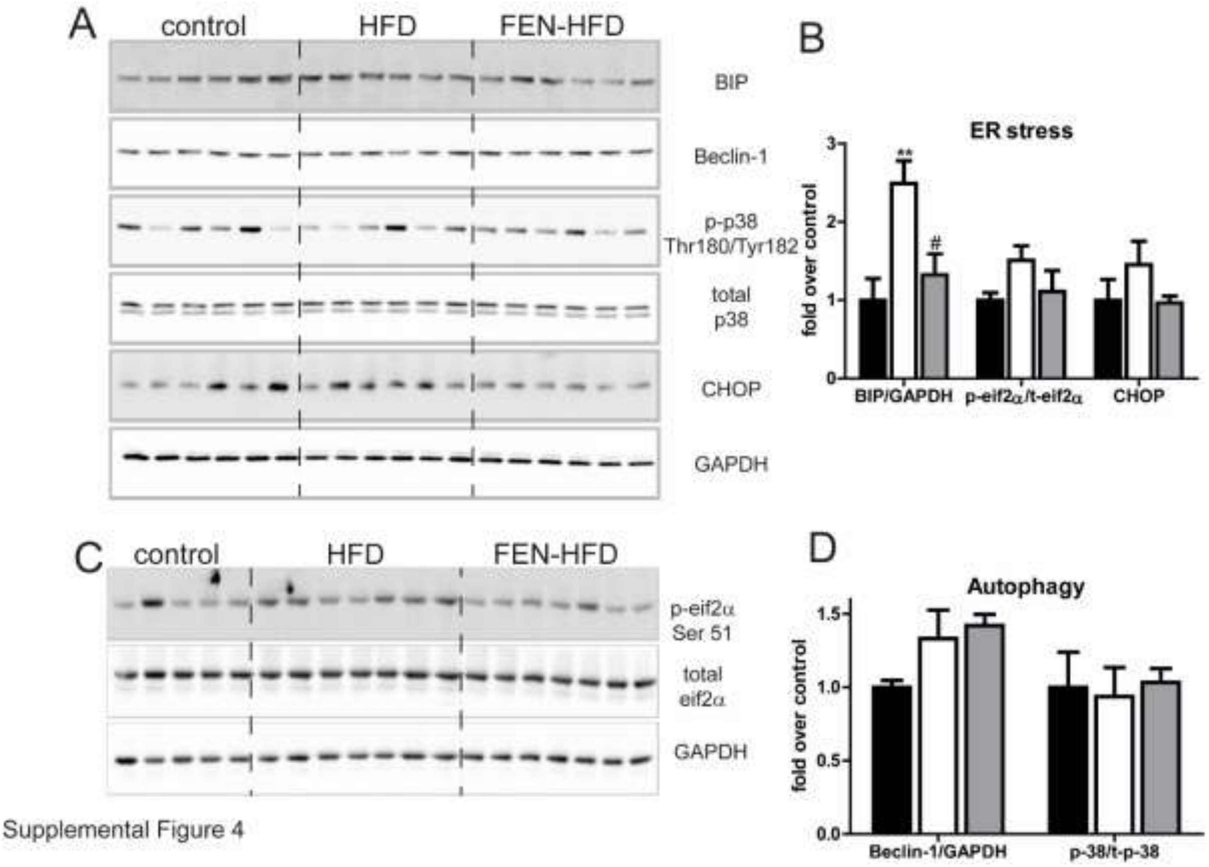


**Supplemental Figure 4: Fenretinide alters hepatic ER Stress but not autophagy.** (**A, C**) Western blot analysis of hepatic tissues probed for BIP, Beclin-1 p-eif2α (Ser51), total eif2α, p-p38 (Thr 180/Tyr 182), total p38 or CHOP. GAPDH was used as a loading control. (**B, D**) Quantification of proteins shown in (**A, B**). Data are represented as mean + S.E.M. and analysed by one-way ANOVA followed by Bonferroni multiple comparison t-tests where $^{**}p \leq 0.01$ (control compared to HFD) or $\#p \leq 0.05$ (HFD compared to FEN-HFD).

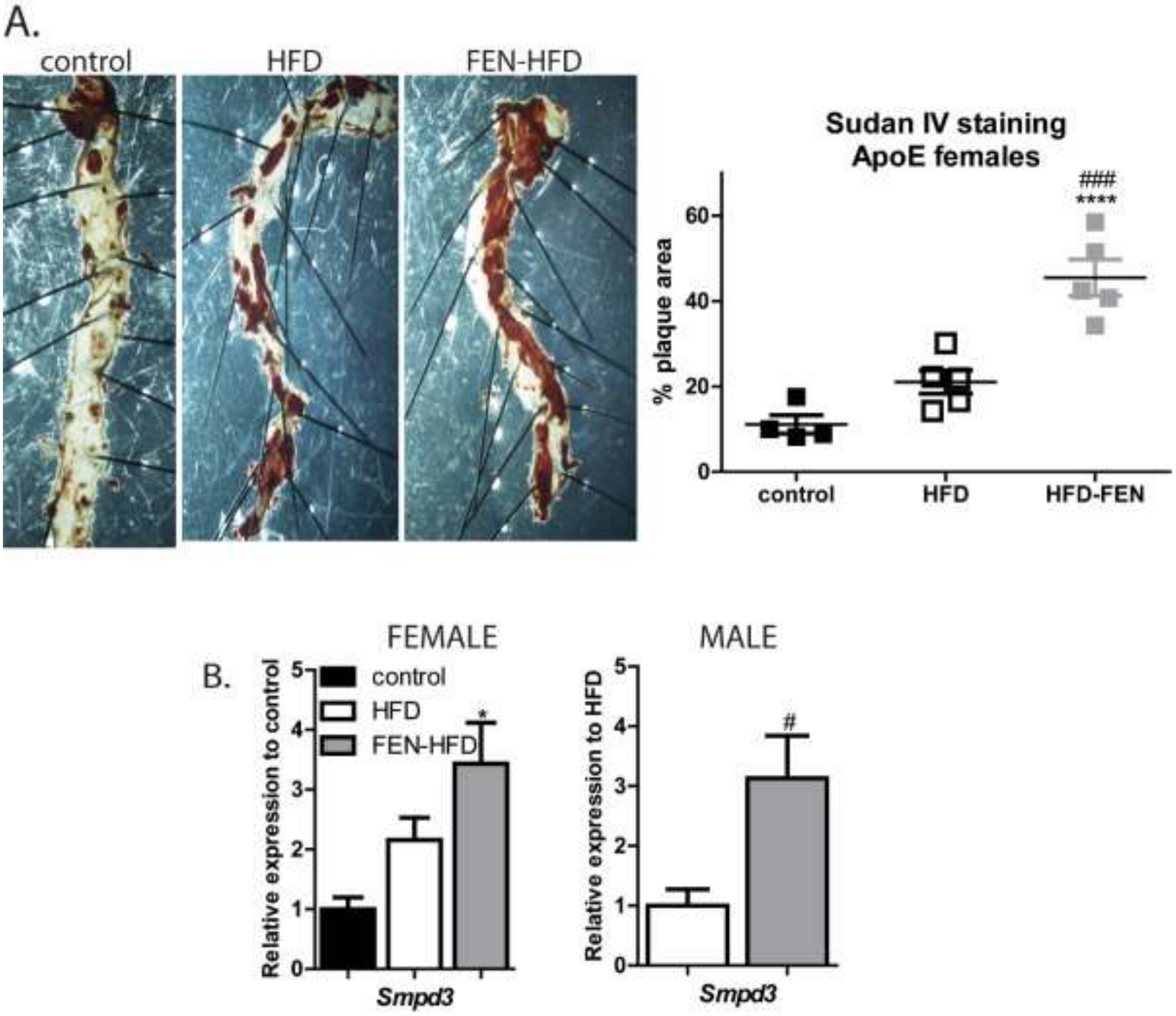


Supplemental Figure 5

**Supplemental Figure 5: Fenretinide increases atherosclerotic plaque formation in female ApoE$^{-/-}$ mice.** The descending aorta was prepared *en face* from (**A**) female ApoE$^{-/-}$ mice and stained with Sudan IV and quantified using Image J. Representative images are shown. Genetic analysis of *Smpd3* levels in hepatic tissue of female (**B**) and male (**C**) ApoE$^{-/-}$ mice. Data are represented as mean ± S.E.M. (n=3-5 per group) and analysed by one-way ANOVA followed by Bonferroni multiple comparison t-tests where *p≤0.05 and ****p≤0.0001 (compared to control) or # p≤0.05 and ### p≤0.001 (HFD compared to HFD-FEN).

A

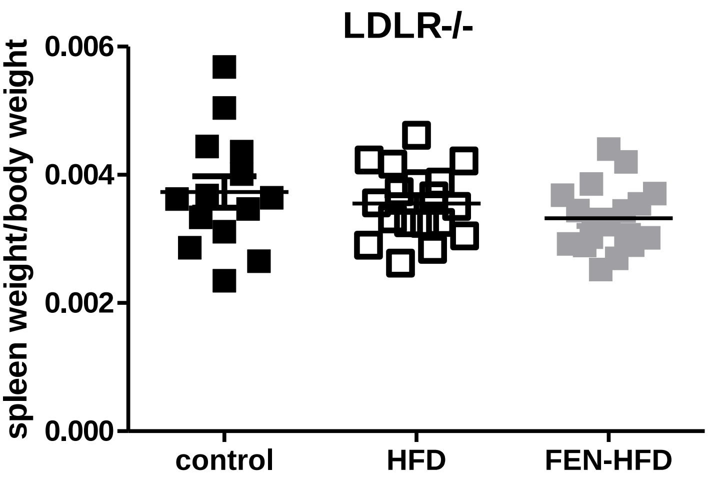


B

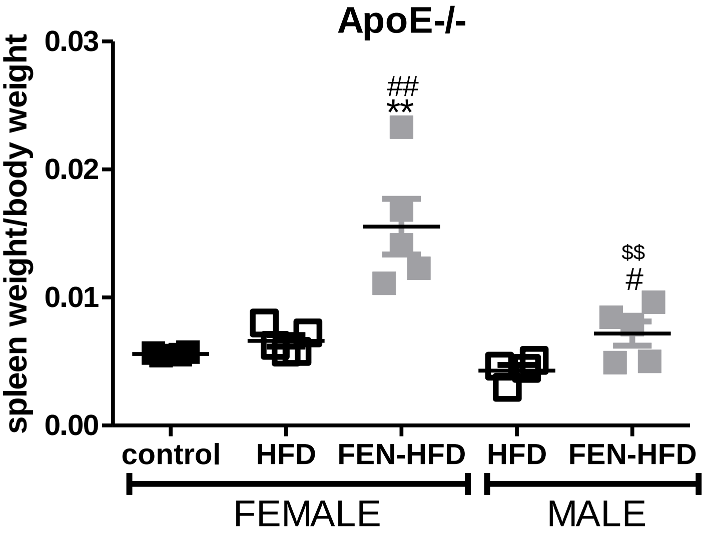


**Supplemental Figure 6: Fenretinide does not induce splenomegaly in LDLR$^{-/-}$ but does in ApoE$^{-/-}$ mice.** Individual spleens were weighed and normalised for body weight of individual mice. Data are represented as mean ± S.E.M. (A. n=14-18 per group LDLR$^{-/-}$ mice; B. n=4-5 per group ApoE$^{-/-}$ mice) and analysed by one-way ANOVA followed by Bonferroni multiple comparison t-tests where **$p \le 0.01$ (compared to control), # $p \le 0.05$ and ## $p \le 0.01$ (HFD compared to HFD-FEN) and $$ $p \le 0.01$ (ApoE$^{-/-}$ FEN-HFD, male compared to female).
